# Analysis of alternative mRNA splicing in vemurafenib-resistant melanoma cells

**DOI:** 10.1101/2022.03.16.484656

**Authors:** Honey Bokharaie, Walter Kolch, Aleksandar Krstic

## Abstract

Alternative mRNA splicing is common in cancers. In BRAF V600E mutated malignant melanoma a frequent mechanism of acquired resistance to BRAF inhibitors involves alternative splicing (AS) of BRAF. The resulting shortened BRAF protein constitutively dimerizes and conveys drug resistance. Here, we have analysed AS in SKMEL-239 melanoma cells and a BRAF inhibitor (vemurafenib) resistant derivative that expresses an AS, shortened BRAF V600E transcript. Transcriptome analysis showed differential expression of spliceosome components between the two cell lines. As there is no consensus approach to analysing AS events, we used and compared four common AS softwares based on different principles, DEXSeq, rMATS, ASpli, and LeafCutter. Two of them correctly identified the BRAF V600E AS in the vemurafenib resistant cells. Only 12 AS events were identified by all four softwares. Testing the AS predictions experimentally showed that these overlapping predictions are highly accurate. Interestingly, they identified AS caused alterations in the expression of melanin synthesis and cell migration genes in the vemurafenib resistant cells. This analysis shows that combining different AS analysis approaches produce reliable results and meaningful, biologically testable hypotheses.

**Graphic Abstract:** 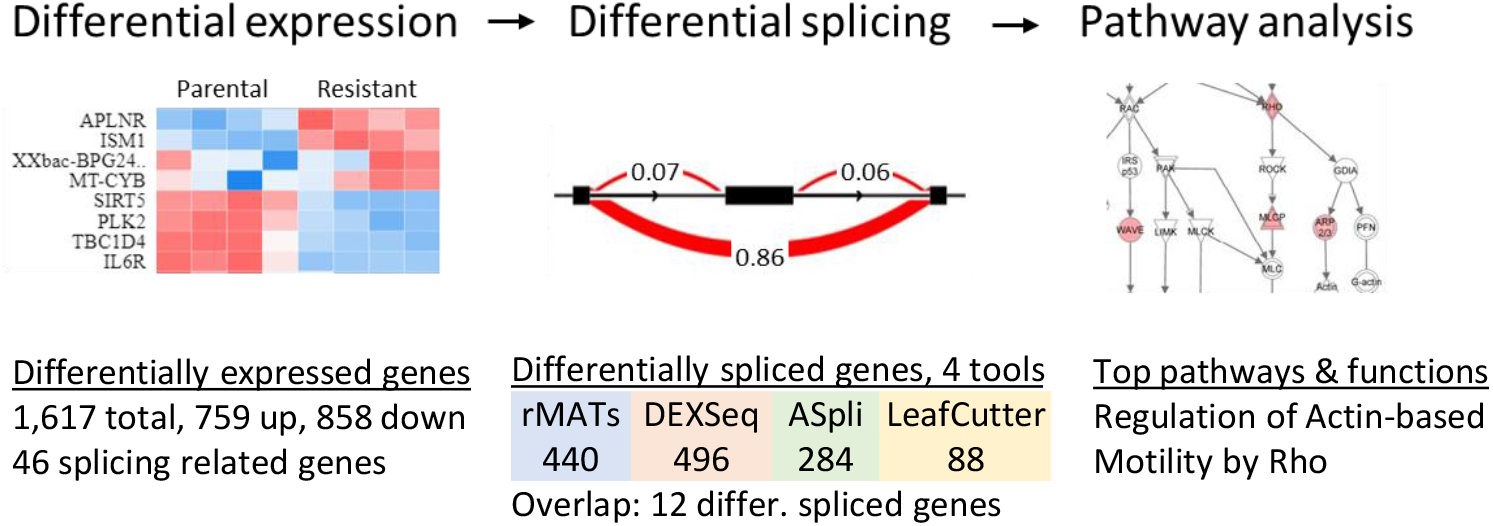

## 1. Introduction

Malignant melanoma is a cancer that originates from melanocytes and is ranked 21st among most common cancers [1, 2]. In 2018, 287,723 new cases of melanoma and 60,712 deaths were registered worldwide. Even though melanoma constitutes less than one percent of skin cancer cases, it is highly malignant and responsible for 79% of skin cancer-related deaths [1, 2].

Several mutations in melanoma activate signaling pathways that regulate cell proliferation. BRAF, NRAS, and NF1 mutations, all activate the MEK/ERK pathway. The MEK/ERK pathway is a signaling cascade that transduces proliferative signals from the extracellular environment to the nucleus of the receiving cells [3]. Normally, the pathway is activated by extracellular ligands, such as growth factors, that bind to receptor tyrosine kinases. These ligand-bound receptors then activate RAS GTPases, which leads to the dimerization and phosphorylation of RAF protein kinases and the subsequent phosphorylation and activation of the MEK and ERK kinases. Activated ERK can stimulate several transcription factors and regulate genes involved in many cellular processes including cell proliferation. In cancer cells the MEK/ERK pathway is often constitutively activated by mutations, thus promoting the oncogenic transformation of the mutated cells [3]. The most frequent type of mutations in melanoma are mutations in the BRAF oncogene (>60% of cases). In 2002 the cancer genome project identified BRAF mutations in more than 60% of melanomas [4]. BRAF is a serine/threonine-protein kinase, and these BRAF mutations constitutively activate BRAF kinase activity and the downstream ERK pathway [4]. Activation of ERK signaling was confirmed as an early event in human melanoma in 2002 by Cohen *et al*. [5]. Among the more than 20 different BRAF mutations in melanoma, the BRAFV600E mutation is the most prevalent and accounts for 90% of all BRAF mutations in melanomas.

Because of the prevalence of the BRAF mutations in melanoma, one of the most successful targeted therapies for BRAF mutated melanomas are BRAF kinase inhibitors such as vemurafenib [6]. Like all targeted inhibitors, vemurafenib suffers from the development of resistance leading to patient relapse. In fact, more than 80% of patients experience relapse within eight months of vemurafenib treatment [7]. Resistance mechanisms of BRAF inhibition are chiefly mediated by ERK pathway reactivation, often by directed BRAF alterations such as BRAF alternative splicing, gene amplification, double kinase fusions and deletions of the BRAF N terminus [8]. Of those, one of the most common mechanisms in melanoma is the alternative splicing (AS) of BRAF, which occurs in 15-30% of patients [9].

AS is one of the molecular hallmarks of human cancer [10]. Cancer has about 30% more AS events than normal tissue, often producing disease-specific protein isoforms [11, 12]. mRNA splicing is mediated by the spliceosome, which is a large complex comprised of five small nuclear ribonucleoproteins U1, U2, U4, U5, U6, and splicing factors including SR proteins, heterogeneous nuclear ribonucleoproteins (hnRNPs), and auxiliary proteins [13]. SR proteins regulate splicing by attaching to exonic and intronic splice enhancer sites, which are sequence motifs within exons and introns [14]. Similarly, hnRNPs regulate splicing by binding to silencer sites that block the access of spliceosome elements and inhibit splicing at these sites [13]. Auxiliary proteins are involved in the assembly of the core-splicing complex to make a functional complex that can produce different splice isoforms from the same gene [13].

Vemurafenib resistant melanoma cells often express an alternatively spliced short BRAFV600E isoforms that lack the RAS-binding domain. In a study of 19 patients that acquired resistance to vemurafenib, four short isoforms were observed in six patients with transcripts lacking exons 4-10, exons 4-8, exons 2-8, or exons 2-10 [15].

A suitable cell model system to study BRAF-splicing mediated vemurafenib resistance are SK-MEL-239 melanoma cells that had acquired resistance [15]. Similar to what was observed in patients, this cell line expresses a short BRAF splice variant that lacks the RAS-binding domain. This splice variant shows enhanced dimerization, which drastically enhances kinase activity [16] [17] [18], thus leading to persistent activation of the RAF/MEK/ERK pathway even in the presence of vemurafenib.

Considering that AS is common in cancer, aberrant splicing of BRAF might not be the only splicing event related to vemurafenib resistance. Hence, we sought to characterise systematically the aberrant splicing landscape in vemurafenib resistant cells.

## 2. Materials and Methods

### 2.1 Cell culture and treatments

SK-MEL-239 clone C3 cell line was received as a generous gift from Prof. Poulikos I. Poulikakos (Department of Oncological Sciences Icahn School of Medicine at Mount Sinai, New York, USA). The establishment of the cell line clone is described by Poulikakos et al. 2011 [15]. The culturing conditions described in the original publication were used. Briefly, parental SK-MEL-239 cells were cultured in RPMI 1640 medium (Gibco) supplemented with 10% FBS (Gibco), penicillin/streptomycin (1X) (Gibco) and L-glutamine (1x) (Gibco). Resistant SK-MEL-239 clone 3 were cultured in the same media supplemented with 2 µM vemurafenib (PLX4032) (SelleckChem). Cells were cultivated in cell culture incubator (Thermo Scientific) at 37°C and 5% CO_2_. Depleted medium was replaced with fresh pre-warmed media every two to three days.

### 2.2 Cell Viability Assay

Relative cell viability was measured by MTS assays using the CellTiter 96^®^ AQueous One Solution Cell Proliferation Assay (Promega). Briefly, SK-MEL-239 cells, parental and vemurafenib resistant clone 3, were seeded in 96-well flat-bottom plates (1 × 10^4^ cells/well) with 100 µL of 10% FCS media and incubated for 24 hours. Graded dilutions of vemurafenib or DMSO vehicle control, in culture medium, were added to each well in triplicate. Upon drug treatment, MTS reagent was added and, after one hour of incubation, the absorbance at 490 nm was measured using the plate reader (Spectramax Plus384 Plate Reader, Molecular Devices accompanied with SoftMaxPro software). The results were background-corrected by subtracting the average signal of wells only containing medium, and normalized to the no treatment control at the corresponding 48 and 72 hours timepoint. The mean ± standard deviation (SD) of triplicate samples, were calculated and plotted against the increasing concentration of vemurafenib treatments (0.078-10 µM) Microsoft Office Excel.

### 2.3 Western Blot

Parental and vemurafenib resistant SK-MEL-239 cells were seeded in 6 well plates and allowed to grow for 24 hours. Then, culture medium was replaced with fresh medium without drugs or with vemurafenib, at 1 µM or 10 µM concentration. After 30 or 60 min, cells were placed on ice, washed with ice-cold 1x PBS and harvested using 600 µl of lysis buffer (5% NP40, 10 mM Tris-HCl (pH 7.5), 150 mM NaCl supplemented with protease inhibitor cocktail (cOmplete™ Mini Protease Inhibitor Cocktail, Roche Diagnostics) and phosphatase inhibitor cocktail (PhosSTOP, Roche Diagnostics).

Lysates were cleared by centrifugation at 12,000 rpm for 20 min at 4°C, transferred to fresh tubes and stored at −20°C. The prepared whole cell lysates were mixed with 4x Loading buffer (44.4% glycerol, 4.4% SDS, 277.8mM Tris pH 6.8 and 0.02% Bromophenol blue, 100 mM DTT), heated for 5 min at 95°C, cooled on ice and resolved using SDS-PAGE electrophoresis. The Precision Plus Protein™ Dual Colour ladder (BioRad) was used as a molecular weight standard. Upon transfer onto PVDF membrane (Hybond-P, Amersham), membranes were blocked in 5% non-fat milk (Sigma) in TBST at room temperature for 1 hour. Membranes were probed with primary antibodies diluted in 5% (w/v) BSA in 1x TBST overnight at +4°C. Next day, membranes were washed and incubated in the horseradish peroxidase (HRP)-conjugated secondary antibodies directed against primary mouse and rabbit antibodies (dilution of 1/5000 in 5% (w/v) non-fat milk powder in1x TBST), for 1 hour at room temperature. Next, membranes were incubated with ECL substrate (Pierce ECL Western Blotting Substrate, Thermo Fisher) and the chemiluminescent signal was acquired with Chemi Imager (Advanced Molecular Vision accompanied with Chemostar software). Primary antibodies directed against following proteins were used: BRAF (F-7) (1/1000 dilution, #sc-5284, Santa Cruz, Mouse), pMEK (1/1000 dilution, #9121, Cell Signaling Technology, Rabbit), ppERK1/2 (E-4) (1/10,000 dilution, #sc-7383, Santa Cruz, Mouse), tERK1/2 (1/10,000 dilution, #M5670, Sigma, Rabbit),pRSK-1/2 (1/1000 dilution, #sc-12898-R, Santa Cruz, Rabbit), SF3B1 / SAP 155 (B-3) (1/1000 dilution, #sc-514655, Santa Cruz, Mouse), HSP90 (c45g5) (1/1000 dilution, #4877, Cell Signalling Technology, Rabbit), GAPDH (14C10) (1/1000 dilution, #2118, Cell Signalling Technology, Rabbit). Following horseradish (HRP)-conjugated peroxidase secondary antibodies were used: Anti-mouse secondary antibody (1/5000 dilution, #7076, Cell Signalling Technology, Horse), Anti-rabbit secondary antibody (1/5000 dilution, #7074, Cell Signalling Technology, Goat).

### 2.4 RNA Sequencing

Total mRNA was extracted from four parental SK-MEL-239 and four vemurafenib resistant SK-MEL-239 RNA biological replicates, using RNeasy Mini Kit (Qiagen) according to manufacturer’s protocol, and DNA was digested with DNA-free™ DNA Removal Kit (Applied Biosystems). RNA integrity was assessed on 2100 Bioanalyzer (Agilent) using a Eukaryote Total RNA Nano Chip (version 2.6), with samples’ RIN value range ranging from 8.9 to 10. Poly A selection was performed using NEB Next® Ultra ™ RNA Library Prep Kit (New England Biolabs) and the sequencing libraries (250∼300 bp insert cDNA library) were generated with a proprietary methodology developed by Novogene, China. 150 bp paired-end sequencing was performed on Illumina NovaSeq 6000 platform (Novogene, China).

### 2.5 PCR

1 µg of the total RNA was reverse transcribed using the QuantiTect Reverse Transcription Kit (Qiagen) according to the manufacturer’s protocol. RT-PCR amplification for the detection of selected splice variants was performed for the following genes: TYR, EPB41, CLSTN1, MPRIP, FANCA, MARK3, EVI5L, CAPN3, BRAF as well as for the housekeeping control gene GAPDH. Exon-exon junction spanning primers were designed using Oligo7 https://www.oligo.net) [19] and optimal design parameters were double checked with Generunner (http://www.generunner.net/) and Primer3Plus (https://pri-mer3plus.com/). Additionally, Primer-BLAST (https://www.ncbi.nlm.nih.gov/tools/primer-blast/) was used to eliminate designed primers with unspecific binding. Primer sequences are provided in the Supplemental Material, Table S6.

PCR reactions were performed in 25 μl reactions using the MyTaq Red Mix (Bioline, Meridian Life Science). The RT-PCR reaction conditions were optimised for each primer pair and are designated as Condition A to D. For Condition A amplification parameters are: denaturation 1 min at 95ºC, followed by 35 cycles of denaturation at 95ºC for 30 s, annealing at 60ºC for 30 s and elongation at 72ºC for 10 s, followed by 10 min elongation at 72ºC. For Condition B amplification parameters are: denaturation 1 min at 95ºC, followed by 30 cycles of denaturation at 95ºC for 30 s, annealing at 59ºC for 30 s and elongation at 72ºC for 10 s, followed by 10 min elongation at 72ºC. For Condition C amplification parameters are: denaturation 2 min at 95ºC, followed by 30 cycles of denaturation at 95ºC for 30 s, annealing at 61ºC for 30 s and elongation at 72ºC for 40 s, followed by 10 min elongation at 72ºC. For Condition D, amplification parameters are: denaturation 1 min at 95ºC, followed by 35 cycles of denaturation at 95ºC for 30 s, 62ºC for 30 s and 72ºC for 10 s, followed by 10 min elongation at 72ºC. For Condition D, amplification parameters are: denaturation 1 min at 95ºC, followed by 35 cycles of denaturation at 95ºC for 15 s, annealing at 56ºC for 15 s and elongation at 72ºC for 10 s, followed by 10 min elongation at 72ºC. RT-PCR products were separated by gel electrophoresis in 1% agarose (Sigma). GeneRuler 100 bp DNA ladder (ThermoScientific) was used as a marker, and digital images of the gels were taken with MiniBis gel doc system (software GelCapture v7.4; DNR Bio-Imaging Systems)

### 2.6 Ingenuity Pathway Analysis

Functional enrichment analysis was performed by using Ingenuity Pathway Analysis (IPA; Qiagen, https://www.qiagenbioinformatics.com/products/ingenuitypathway-analysis). First, differentially expressed genes were uploaded to IPA and a pathway enrichment analysis against the Ingenuity Knowledge Base was performed. To identify how many splicing genes were differentially expressed, a BioProfiler analysis for the GO biological function “alternative splicing” was performed. The IPA BioProfiler analysis probes the repository of scientific information to generate molecular profiles of diseases, phenotypes and biological processes (e.g., alternative splicing) listing all the genes and compounds that have been associated with the profiled term. Also, the lists of differentially spliced genes from each splicing analysis tool were uploaded to IPA. The pathway enrichment analysis was performed separately for each tool, then a comparative analysis was performed, for which the tools were treated as multiple conditions. For all analyses, p-values for pathway over-representation analysis were generated by IPA using a right-sided Fisher exact test and Benjamini–Hochberg correction for multiple hypothesis testing. p-values < 0.05 were considered significant.

### 2.7 Galaxy platform

The sequencing quality control analysis was performed on the Galaxy web platform (usegalaxy.org) [20] using FastQC to obtain phred scores for assessing base-calling accuracy and GC content [21]. The paired-end sequence reads (FASTQ files) were aligned to the human reference genome GRCh38 (hg38, GenBank assembly accession: GCA_000001405.28) using HISAT2 aligner [22], also on the Galaxy public server. Alignment files were used for the downstream AS analyses.

### 2.8 Biojupies

Differential gene expression analysis was performed on BioJupies web platform (https://amp.pharm.mssm.edu/biojupies/) [23]. The analysis was performed using tools for Principal Component Analysis (PCA), gene clustering (Clustergrammer), differential expression analysis and volcano plot diagrams. All diagrams are generated using the embedded Plotly tool (https://plot.ly).

### 2.9 Differential splicing analysis

Differential splicing analysis was performed using four different tools.

ASpli (Version 2.0.0) is as part of the Bioconductor R package (Release 3.12, DOI: 10.18129/B9.bioc.ASpli), and it makes use of junction reads information and quantifies the pre-mRNA splicing events through calculating PSI and PIR matrix [24]. The AS events with an absolute FDR < 5% and Delta PSI_PIR > 3% were deemed differentially spliced.

DEXSeq (Version1.36.0) is a part of the Bioconductor R package (Release 3.12, DOI: 10.18129/B9.bioc.DEXSeq), and it identifies AS through inferring the relative exon usage within each gene [25]. Cut-offs: FDR<0.05, logFC >2.

LeafCutter (Version 0.29) was obtained from GitHub (https://github.com/davidaknowles/LeafCutter) and was installed *via* the R devtools package devtools::install_github (“davidaknowles/LeafCutter/LeafCutter”) [26]. This package identifies AS events by intron-based clustering approach, where splicing is measured as the excision of introns.

Two packages, rMATS (Version 4.1.0) for differential splicing and Maser (Version 1.7.0), were used for annotating the splicing events with protein domains. rMATS was obtained from the open-source platform SourceForge (http://RNA-seq-mats.source-forge.net/). Maser was obtained from Bioconductor (Release 3.112) DOI: 10.18129/B9.bioc.maser). These two tools are based on quantifying and annotation of exon-included and exon-excluded junction-spanning reads for each AS event [27].

Scripts for all four AS tools are provided in Supplemental Data 1.

## 3. Results

### 3.1 Characterization of the cell line model system

For this study we used the human cell-line model system for acquired vemurafenib resistance in malignant melanoma established by Poulikakos et al. [15]. This model system consists of a pair of isogenic human melanoma cells with a BRAF V600E mutation, i.e., parental and drug resistant SK-MEL-239 cells. Vemurafenib resistant clones were generated from parental SK-MEL-239 cells through continuous long term drug exposure. Here, we used the resistant clone C3, which expresses a short BRAF splice isoform of 61 kDa in addition to the full length BRAF isoform of 85 kDa. To assure that the parental cells and Clone 3 respond differentially to vemurafenib, we treated them with 1 µm and 10 µM vemurafenib for 30 and 60 minutes and measured the phosphorylation of MEK, ERK, and the ERK substrate RSK1/2 (Figure S1). In line with the original report by Poulikakos et al. [15] we observed that parental SK-MEL-239 cells express the full length 85 kDa BRAF isoform (p85), while the resistant clone C3 expressed both the p85 full length and the alternatively spliced 61 kDa BRAF isoform (p61) (Figure S1). ERK signalling, as assessed by monitoring activating phosphorylation sites in MEK, ERK and RSK, was completely inhibited by vemurafenib in parental cells under all conditions. By contrast, in resistant cells there was no inhibition with 1 µM of vemurafenib and only partial inhibition with 10 µM vemurafenib (Figure S2). These results confirmed that the model system has the same characteristics as described in the original report by Poulikakos et al. [15].

To measure the dose- and time-dependent effects of vemurafenib on the viability of SK-MEL-239 cells, we used the MTS cell viability assay (Figure S2). In parental cells, 10 µM vemurafenib reduced cell viability to 52% and 28% after 48 and 72 hours of treatment, respectively (Figure S2). In resistant cells, no marked differences of cell viability were observed for any of the drug doses and length of treatment times. These experiments confirmed that parental SK-MEL-293 cells were sensitive to vemurafenib, while the C3 cells were resistant even beyond the 2 μM vemurafenib dose that was routinely included in the growth medium [15].

### 3.2 11 spliceosome genes are differentially expressed in resistant cells

To investigate any changes in transcriptional and AS landscape caused by drug resistance in this model system, we performed RNA-seq in parental and clone C3 SK-MEL-239 cells. RNA-seq data were analysed on BioJupies (https://maayanlab.cloud/biojupies/) and Galaxy (https://usegalaxy.eu/) servers, which enabled us to perform customized analysis with well-established and state of the art RNA-seq pipelines [20, 23]. The FastQC quality control analysis [21] on the Galaxy platform showed at least 30 million 150 bp paired-end reads per each of the four biological replicates, with the average phred score of more than 35 across all base pair positions, unbiased and normally distributed GC, confirming the high quality and deep coverage of the RNA-seq data that is important for a reliable AS analysis. Principal Component Analysis (PCA) showed that 87% of the variance in the RNA-seq data was explained by the first principal component, clearly distinguishing parental and resistant SK-MEL-239 cells (Figure S3). 1617 genes were differentially expressed (759 genes were upregulated and 858 were downregulated) at cut-offs of adjusted p-value<0.05 and log2 fold-change>1 (Table S1). Clustering was performed for the top 50 variable genes. The result shows two strong clusters of upregulated and down-regulated genes that clearly distinguish the parental from the resistant cells (Figure S4).

Because AS of BRAF is the mechanism of resistance in these cells, we examined the genes related to splicing and the spliceosome. First, we looked at all the genes with the GO term “RNA Splicing” (478 genes) and called a gene differentially expressed when the FDR was < 0.05 and the absolute log2 fold change was > 0.5. Results showed that 46 genes were differentially expressed (Table S2). Next, we examined the genes that form the spliceosome as defined by the Molecular Signature Database (KEGG_SPLICEOSOME, M2044, 127 genes). Applying the same cut-off criteria, this analysis revealed that 11 spliceosome genes were differentially expressed, nine were downregulated and two were upregulated (Figure 1A). The two most differentially expressed genes were SNRPD3, a small nuclear riboprotein (Sm) and DHX38 (PRP16), a helicase that participates in the second step in pre-mRNA splicing. Both genes were highly expressed in parental cells and were one log2 fold-change downregulated in resistant cells. Furthermore, three splicing factors SF3A1, SF3B2, and SF3B3, which belong to U2 complex, were downregulated in resistant cells, whereas two genes, RBM8A (Y14) and BCAS2 (SPF27), were upregulated (Figure 1B).

**Figure 1.**
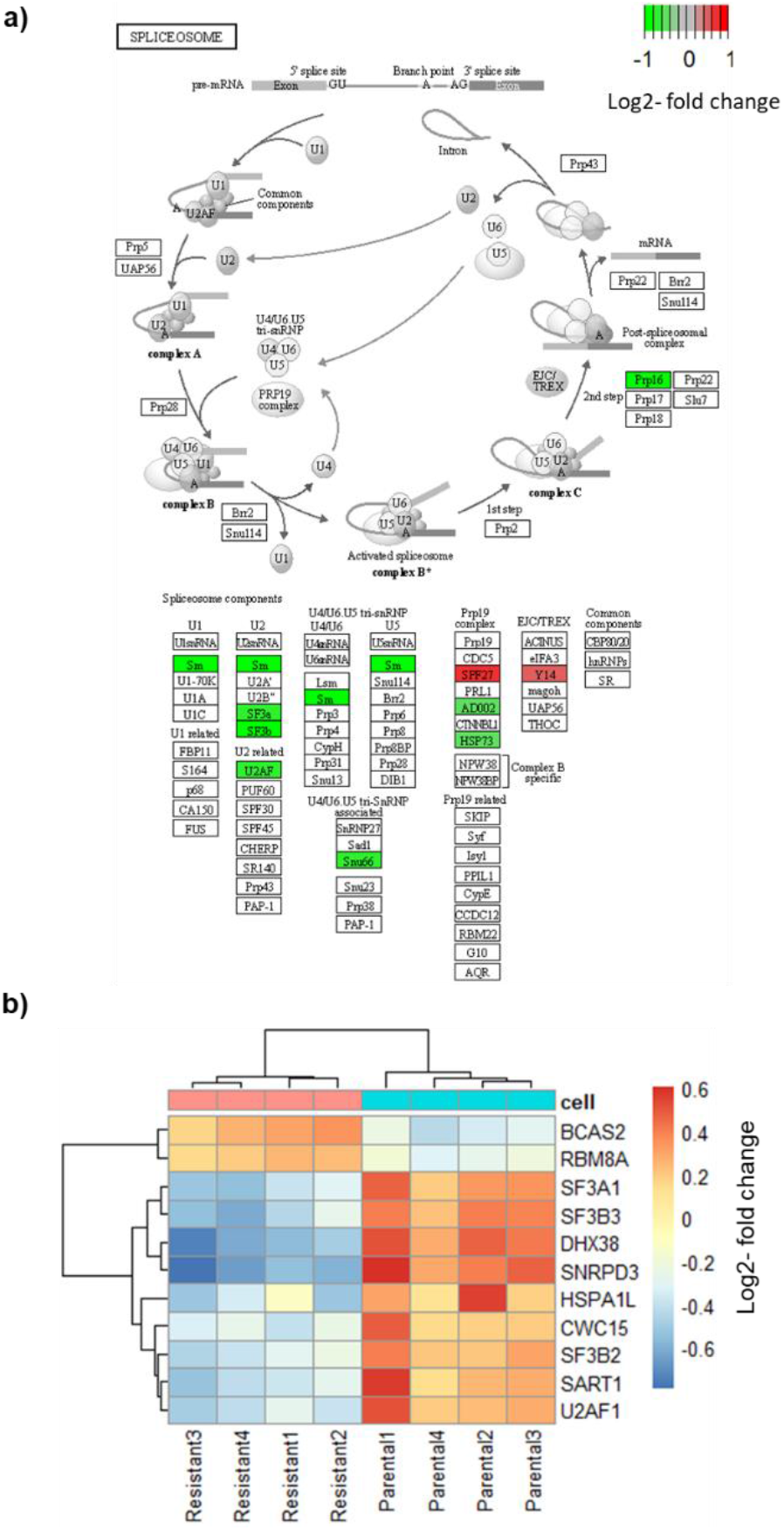
Differential expression of spliceosome genes. a) Scheme of the spliceosome pathway from KEGG (KEGG_SPLICEOSOME, M2044). Red indicates downregulated genes, green upregulated gene. b) Heatmap of differentially expressed spliceosome genes.

### 3.2 Resistant cells exhibit widespread changes in AS

A consensus on approaches for the differential splicing analysis of RNA sequencing data has not been established yet, with common tools differing substantially in their conceptual approach, statistical analyses, and hence in their output data. Therefore, to perform the differential splicing analysis we have employed four different tools that represent three methodological categories: event-based (rMATS-maser [27], LeafCutter [26]), exon usage (DEXSeq [25]), and mixed exon usage and event-based (ASpli [24]) (Figure 2A). The analysis of our RNA-seq data confirmed these observations (Figure 2B). All four tools were used with the same statistical significance cut-offs and identified hundreds of differential splicing events (Figure 2B). DEXSeq identified the most events with 1426, followed by rMATS with 646, ASpli with 544, and LeafCutter with 124 splicing events. The number of differentially spliced genes detected by DEXSeq, rMATS, ASpli, and LeafCutter were 440, 496, 284 and 88, respectively (Figure 2B). All tools call differential exon usage, whereas ASpli and rMATS also call the type of the splicing event such as exon skipping and alternative splice sites usage (Table S3). Exon skipping was the most common type of event (646 of 905 for rMATS).

**Figure 2.**
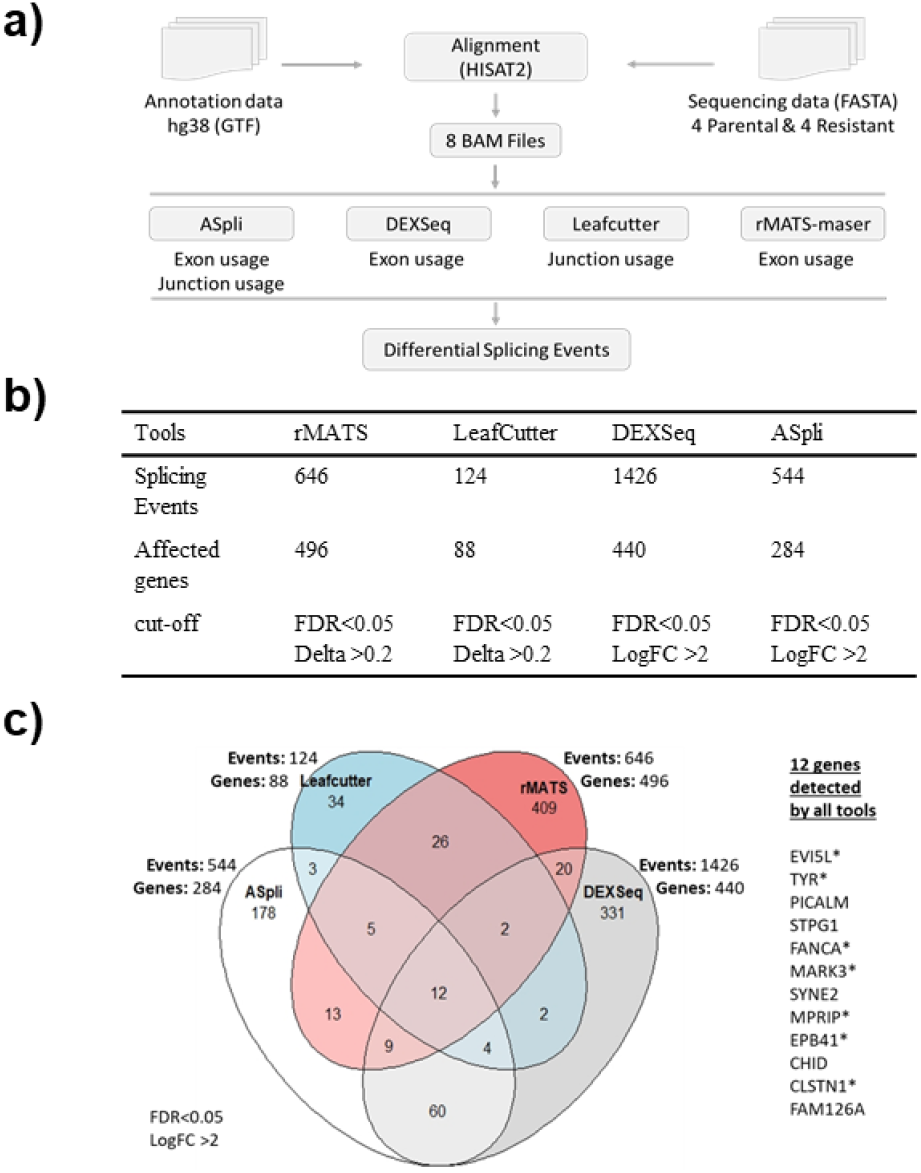
Results of four differential splicing analyses. a) Workflow of the analysis with the four different tools. b) Table showing the number of detected splicing events, affected genes, and cutoffs used for each tool. c) Venn diagram showing the number of splicing events and affected genes for each tool and list of the 12 genes detected by all tools. * indicates the genes selected for further validation.

Thus, we focused on differential exon usage to compare all the tools. As a control for the accuracy of the four softwares, we assessed the AS of the BRAF gene that produces the p61 splice form in SK-MEL-239 C3 cells. ASpli and DEXSeq correctly detected the skipping of BRAF exons 4-8 in C3 cells (Figure 3, Table 1) as originally reported by Poulikakos et al. [15], whereas rMATS and LeafCutter did not detect BRAF splicing (Table S4).

**Table 1.**
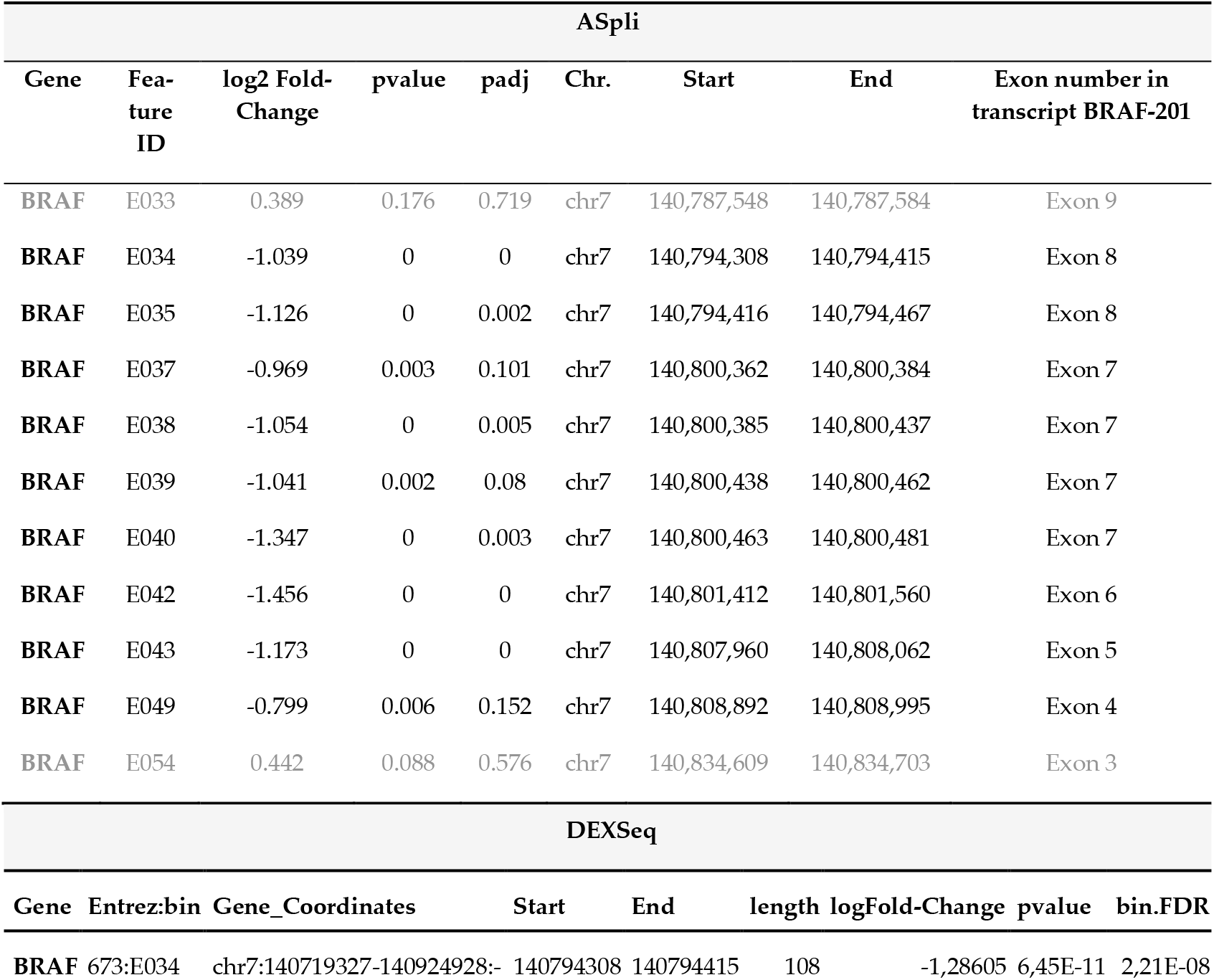
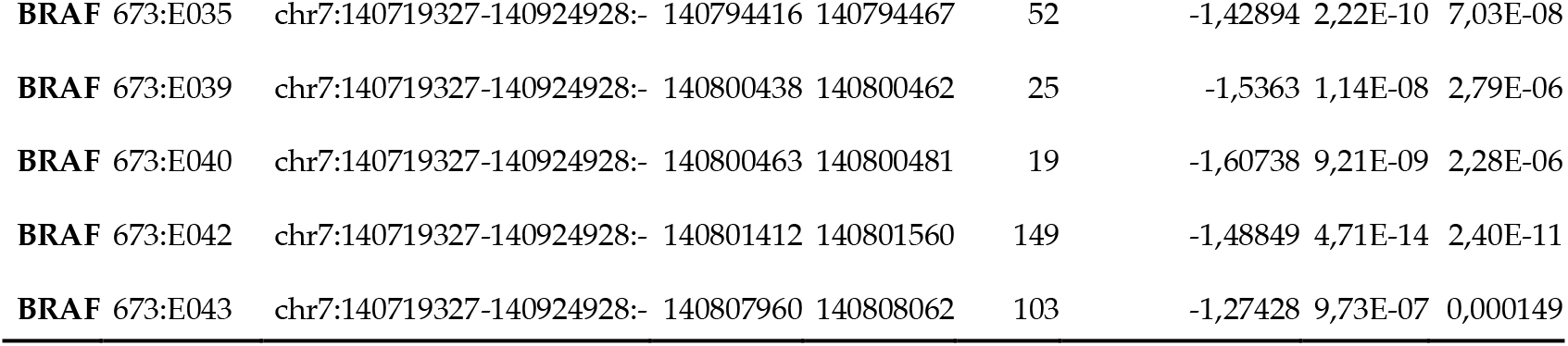
BRAF differential exon usage results from ASpli and DEXseq.

**Figure 3.**
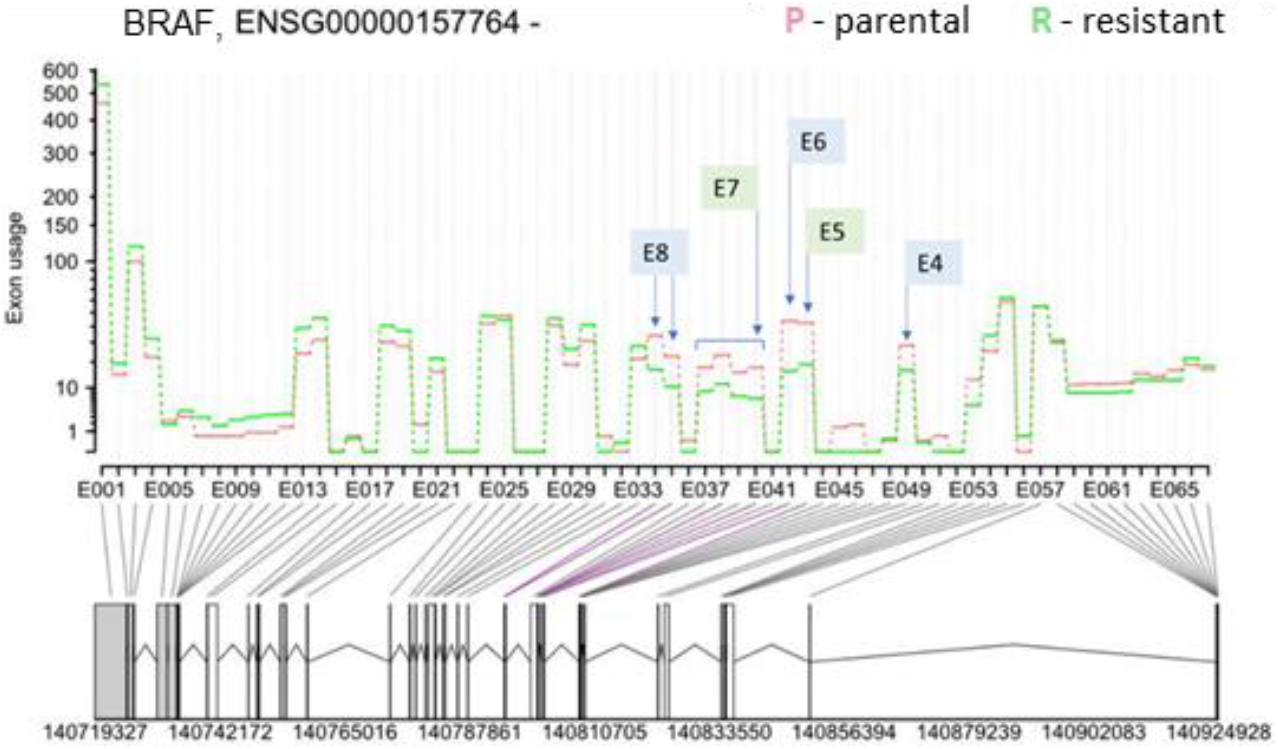
Differential exon usage result for BRAF (Ensembl gene ID: ENSG000000157764) from DEX-Seq. The y-axis shows the exon usage (normalized counts corrected for gene-expression) in parental (P) and resistant (R) cells for each exon bin (x-axis). Exon bins (E001, …) are sections of the genome that correspond to exons in the gene model, as indicated by the grey lines. Below the x-axis the gene model is shown with numbers indicating genomic coordinates. Boxes represent exons. Horizontal lines represent introns. The vertical or diagonal lines indicate the position of the exon bins in the gene model. Purple lines indicate statistically significant bin usage. The positions of the skipped exons 4 to 7 in the resistant cells are indicated by the arrows.

Apart from BRAF, twelve differentially spliced genes were detected by all four tools suggesting that they are bona fide AS events (Figure 2C). Therefore, we analysed the results for the 12 genes from all four tools in detail. First, we compared the genomic locations of the detected splice junctions. For 11 genes (EVI5L, TYR, PICALM, FANCA, MARK3, SYNE2, MPRIP, EPB41, CLSTN1, FAM126A, CAPN3) the same splicing events were detected by all four tools. For CHID1 all four tools detected different events (Table S5).

From the 11 genes detected by all four tools, we chose 7 genes for experimental validation of bioinformatically identified alternative splicing events based on their potential association with melanoma and cancer. We also included CAPN3, although AS was only detected by three tools, because of its association with cisplatin resistance and melanoma aggressiveness [28]. For the experimental AS analysis, we designed primers for sequences in the upstream and downstream exons of the exon that was skipped (Table S6). Expected PCR products would differ in size depending on whether the exon was retained or not (Figure 4). For example, for TYR we designed two pairs of primers. For both pairs of primers, we detected the long PCR product only in the parental cells and the short PCR product only in the resistant cells (Figure 4). The experimental results fully validated the bioinformatics analysis confirming that drug resistance of SK-MEL-239 C3 cells is accompanied by specific AS events.

**Figure 4.**
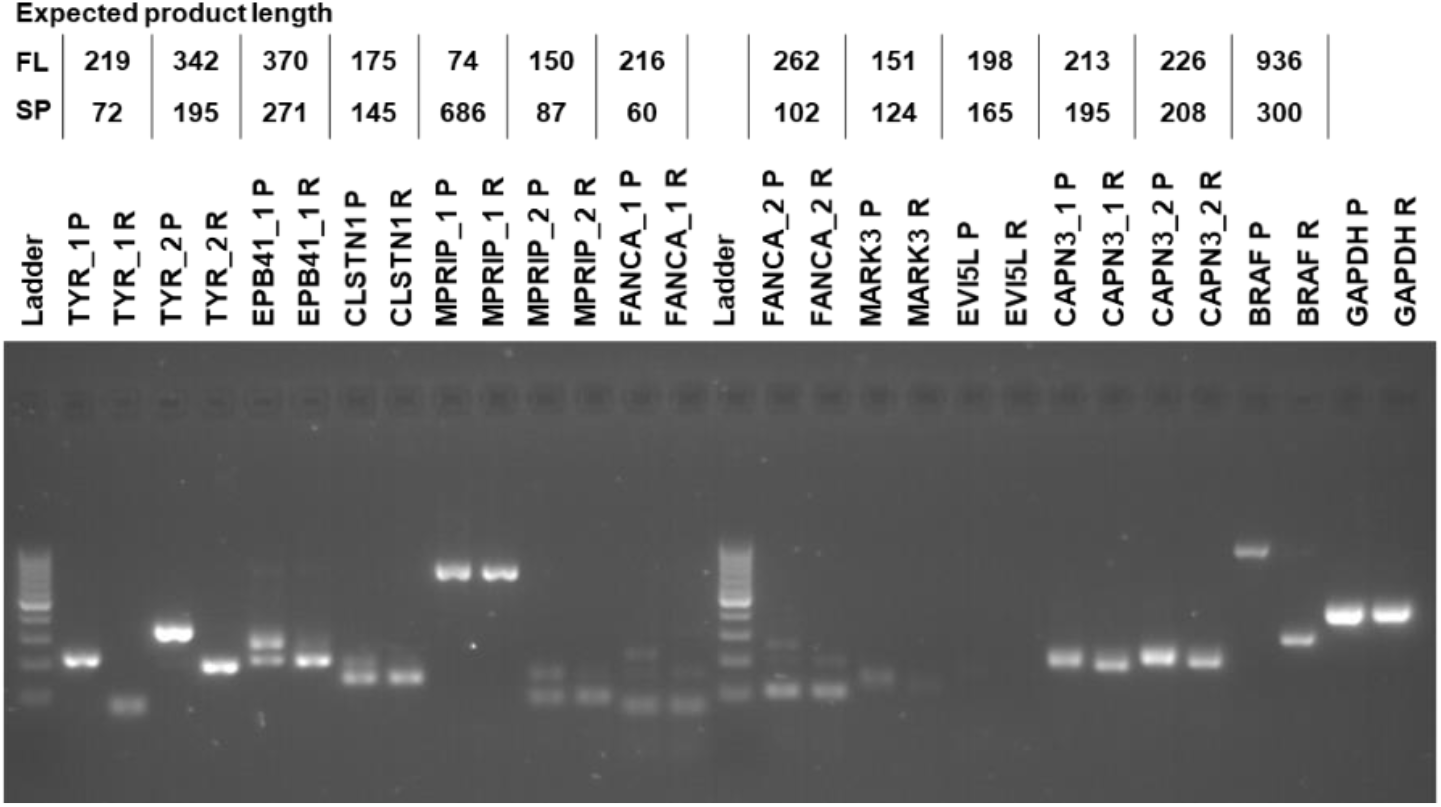
RT-PCR validation. Labels on the top of the gel image provide gene name, primer number in cases when more than one pair of primers were used, parental (P) or resistant (R) sample. Table on top provides the expected RT-PCR product sizes for the full-length (FL) and alternatively spliced (SP) product. Molecular weight DNA ladder marks product sizes from bottom to top: 100, 200, 300, … to1000 bp.

One of the most interesting genes that was alternatively spliced is Tyrosinase (TYR), which is an essential enzyme in melanin synthesis [29]. DEXSeq analysis revealed that a TYR exon located on chromosome 11 from 89,227,823 to 89,227,970 (148bp) was skipped in resistant SK-MEL-239 C3 cells (Figure 5). The skipped exon showed >4000-fold reduction in C3 cells, suggesting an almost complete loss of TYR mRNA containing this exon. To deduce the functional consequence of this splicing event, we inspected the data for an overlap between functional protein domains and this splicing event. For this, we used the Ensembl genome browser, which shows functional protein domains and their locations by using functional domain annotations from databases such as pfam, prints, superfamily and PROSITE (pfam.xfam.org, supfam.org, prosite.expasy.org, respectively). The Ensemble analysis showed two copper-binding domains in the TYR gene (Supplemental Figure S5). Although domain annotations are somewhat different for pfam, superfamily and PROSITE databases, the second copper-binding domain partially or fully overlapped with the spliced-out exon. TYR needs copper binding to function and the spliced-out domain results in the loss of TYR catalytic function [30]. This suggest that vemurafenib resistance is accompanied by AS changes that incapacitate melanin synthesis.

**Fig. 5.**
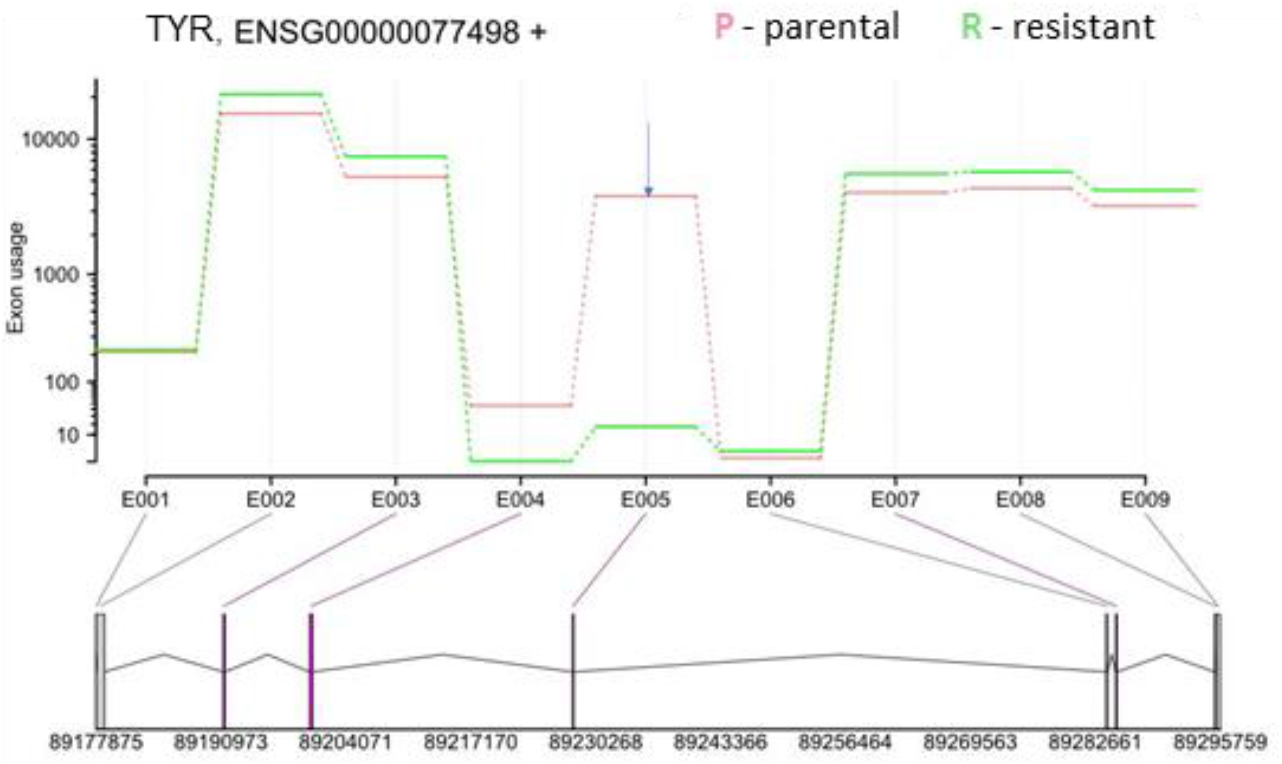
DEXSeq results for TYR (Ensembl gene ID: ENSG00000077498). The y-axis shows the exon usage in parental (P) and resistant (R) cells for each exon bin (x-axis). Below x-axis the gene model is shown with numbers indicating genomic coordinates. Boxes represent exons. Horizontal lines represent introns. The vertical or diagonal lines indicate the position of the bins in the gene model. Purple lines indicate significant bin usage differences.

### 3.3 AS events are correlated with Rho-mediated cell motility

To test whether the alternatively spliced transcripts belong to common pathways, we performed IPA analysis on the results for each AS tool and compared the results. The pathway enrichment analysis detected the “Regulation of Actin-based motility by Rho” pathway as a common pathway for alternatively spliced transcripts identified by all four tools. MPRIP, is the only alternatively spliced gene which is detected by all four tools (Figure 6) (Table S7). An exon on chr17 from 17,180,607 to 17,180,669 of length 63 bp is skipped in resistant cells. In LeafCutter the junction usage for skipping this exon was 0.368 for parental and 0.42 in resistant cells resulting in a delta PSI 0.368 (Figure S5). MPRIP links Rho signalling to actomyosin contractility [31]. The finding of the “Regulation of actin-based motility by Rho” as a common pathway recognized as enriched in the results of four tools (Table S7) is noteworthy, considering that there was limited overlap in the detected differentially spliced genes between the four tools (Table S5). Apart from the AS of MPRIP, which was detected by all four tools, different tools identified different alternatively spliced genes in the pathway (Figure 6).

**Figure 6.**
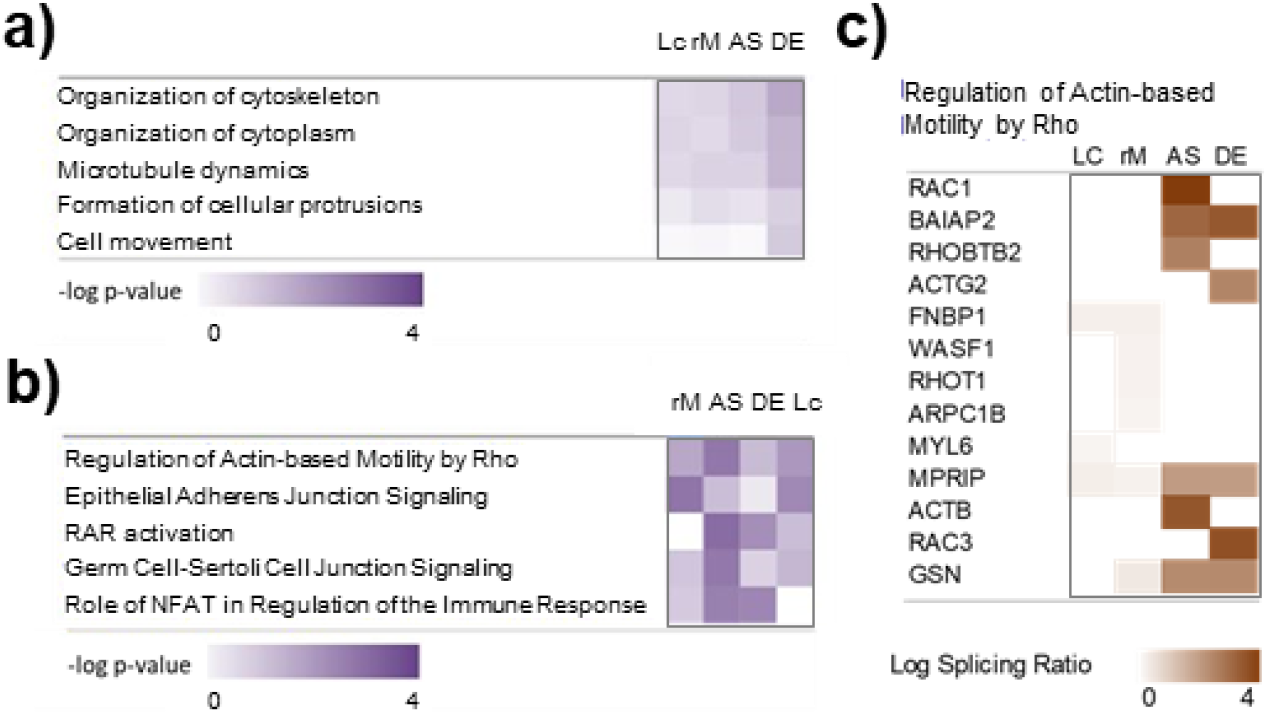
Ingenuity Pathway Analysis (IPA) of the alternative spliced genes. (A) Enrichment of cellular functions. (B) Enrichment of canonical pathways. (C) Heatmap showing which genes were detected as differentially spliced by the different tools in the Actin-based motility by Rho pathway. Lc: LeafCutter; rM: rMATS; AS: ASpli; DE: DEXSeq. Log Splicing Ratio are absolute values of log ΔPSI for rMATS and LeafCutter and log fold change of exon usage for ASpli and DEXSeq.

Rho pathway regulating actin-based motility is well known as an important regulator of cancer invasion and metastasis [32-34] and has also been linked to BRAF inhibitor resistance in melanoma [33, 35]. In the Rho pathway, the Rho-family of GTPases (RhoA, RhoB and RhoC) function as signalling switches that control myosin-actin dynamics[33]. Rho GTPases can switch from an inactive GDP-bound form to an active GTP-bound form. When active, Rho phosphorylates its target Rho-kinase (ROCK). ROCK then controls myosin light chain (MLC) phosphorylation and activity in two ways. Firstly, ROCK directly phosphorylates Myosin Light Chain (MLC), which controls myosin-actin interactions, stress-fibre contraction and cell motility dynamics [36]. Also, ROCK deactivates MLC phosphatase which normally dephosphorylates MLC [37]. Both lead to increased MLC phosphorylation and activity. In this way, activation of the Rho signalling can cause BRAF-inhibitor resistance and was described as a hallmark of therapy resistance in melanoma [33, 35].

## 4. Discussion

For identifying the AS events, we have analysed the RNA sequencing data using four different bioinformatics tools. Each tool identified hundreds of AS events, but only 11 splicing events were in common for all four tools (Figure 2, Table S5). This might be explained by the different identification methods used by each tool. ASpli is a R package specifically designed to deal with the possible complexity of splicing patterns, and considers both bin-based signals and junction inclusion indexes, and uses a generalized linear model [24]. Bins are sequences of the genome split into non-overlapping features. Junctions are features connecting one splice-site to another. DEXSeq is also available as a R package and uses a generalized linear model and uses bins to test for differential exon usage and control false-discovery rates [25]. LeafCutter requires SAMtools, Python and R, but avoids the need of transcript annotations and identifies splicing events from short read RNA sequencing data using a junction-based approach [26]. This approach circumvents the challenges in transcript or exon usage estimation. rMATS requires python and uses a hierarchical model to simultaneously account for sampling uncertainty in individual replicates and variability among replicates and estimates differential exon usage [27].

The differences in our splicing results shows that there is no consensus yet for the analysis of differential splicing. It is well known that different tools use different approaches and therefore recognize different splicing events [38]. But how many of these splicing events are false positives is not clear. The implications are twofold. On the one hand, it could mean that the sensitivity of these tools is limited and that different tools recognize different splicing events. On the other hand, it could mean that these tools suffer from false positives. In this case, using several tools and looking at the overlap will reduce the risk of false positives [38]. This was the approach that we have chosen. The result that all seven tested genes could be validated using PCR shows that using several splicing analysis tools and focusing on the overlap is indeed a good approach for minimising false positives. However, it is possible that many of the other identified splicing events are real. For example, LeafCutter does not require genome annotations in terms of known exons, introns and splice sites, and can thus identifies splicing events that cannot be recognized by the other tools [26].

In line with published results, our differential expression analysis widespread gene expression changes that distinguish resistant cells [39, 40]. A recent study performed RNA-seq analysis in sensitive and resistant A375 melanoma cells found hundreds of differentially expressed genes, but did not analyse alternative splicing [40]. Here, we found differential expression of several splicing factors, including factors of the U2 complex (Figure 1).

Out of the 12 alternatively spliced genes identified by all tools, we have focused our attention on the genes with a putative involvement in the transformation and promotion of the malignant melanoma phenotype. In the following we discuss each of the validated AS events.

One of the alternatively spliced genes with the largest effect size was TYR (Figure 4, Table S3). Both, TYR and its binding protein TYRP1 were also top hits for differential expression. Interestingly, while TYR expression was slightly downregulated, TYRP1, which is involved in the stabilization of TYR, was upregulated in resistant cells, perhaps as a response to the TYR splicing. TYR produces the pigment melanin [29]. Our finding shows that AS of TYR in resistant cells causes the loss of the second copper binding domain by exon skipping (Figure 4, Table S3). The two copper binding domains are important for the TYR catalytic function, suggesting a reduction in melanin pigmentation in resistant cells. A previous study showed TYR downregulation and reduced melanin content in vemurafenib-resistant cells consistent with melanoma cell de-differentiation [41]. Similarly, our result suggests reduced TYR activity resulting from AS as a novel mechanism of TYR de-activation in vemurafenib resistant cells.

CAPN3 (Calpain 3) AS resulted in the loss of exon 15 in resistant cells (Table S3). The expression of two alternatively spliced short isoforms of CAPN3 has been observed before in melanoma, and the downregulation of these isoforms has been linked to melanoma aggressiveness and cisplatin resistance [28]. Both of these short CAPN3 isoforms have exon 15 that contains a nuclear localization signal [28]. The forced expression of these isoforms induced p53 stabilization and cell death in A375 human melanoma cells suggesting that exon 15 is important for the proapoptotic function of CAPN3 [42]. Skipping of exon 15 would mean a loss of the nuclear localization signal and the proapoptotic function of CAPN3. But because the function of exon 15 is not entirely clear [42], this should be tested in future experiments.

Splicing of CLSTN1 (Calsyntenin 1) has previously been recognized as very important in tumour invasiveness [43]. Like in many other cancers, the metastatic process of invasive melanoma is driven by the epithelial–mesenchymal transition (EMT), which is characterized by a loss of E-cadherin and a gain of N-cadherin expression. Whereas the expression of E-cadherin (CDH1) was not altered in the resistant cells, N-cadherin expression was slightly upregulated (log fold-change of 0.3, adjusted p-value 0.018, Table S1). In malignant melanoma, EMT enables melanoma cells to cross the basement membrane of the epidermis into the dermis, which is a critical step in the formation of metastases [44]. A CLSTN1 short isoform has been found to inhibit EMT in breast cancer cells [43]. This short isoform lacks exon 11 of the canonical sequence (Ensembl - transcript CLSTN1-201, ENST00000361311.4). Here, we identified a short isoform in resistant cells that lacks both exon 11, and exon 3. The findings in breast cancer cells suggest that a lack of exon 11 produces a more epithelial phenotype that is less invasive in the resistant cells. Alternatively, this AS event may enhance the reversion of EMT, which is necessary for cells to proliferate once they have settled into a metastatic site [45]. However, our resistant cells also lack exon three, and the biological effects of this are not known. Thus, it would be interesting for future work to determine the effects of the here detected splicing events of CLSTN1 on EMT in melanoma cells.

FANCA (Fanconi Anemia Complementation Group A) is a protein that is involved the Fanconi anemia pathway that is activated when DNA replication is blocked due to DNA damage [46]. Germline coding variations and single-nucleotide polymorphism of the FANCA gene have been associated with melanoma susceptibility [47] and overall patient survival [48], respectively. Our result that FANCA is alternatively spliced suggests alterations of the DNA damage response and repair in resistant cells.

Of the validated alternatively spliced gens, three genes have not yet been associated with melanoma or vemurafenib resistance.

EPB41 (Erythrocyte Membrane Protein Band 4.1), together with spectrin and actin constitutes the cell membrane cytoskeletal network, and plays a key role in regulating membrane mechanical stability and deformability by stabilizing spectrin-actin interaction. The spectrin–actin binding (SAB) domain partially overlaps with the spliced-out exon (ENSE00001065029, exon number 15 in EPB41-201), suggesting that exon skipping results in loss of the EBP41-SAB domain, compromised actin and spectrin binding and destabilization of the cytoskeletal network [49].

MARK3 (Microtubule Affinity Regulating Kinase 3) is a serine/threonine-protein kinase that phosphorylates the microtubule-associated proteins MAP2, MAP4 and MAPT/TAU [50], negatively regulates the Hippo signaling pathway and cooperates with DLG5 to inhibit the kinase activity of STK3/MST2 toward LATS1 [51]. No known protein domain was associated with the skipped exon (ENSE00003477170, exon number 16 in MARK3-205) making it difficult to speculate about the functional consequence.

EVI5L (Ectopic viral integration site 5 like) is a GTPase Activating Protein (GAP) that modulates cell cycle progression, cytokinesis, and cellular membrane traffic [52]. The functional consequence of the skipped exon (ENSE00002211040, exon number 12 in EVI5L-202) is unknown.

The question of what causes the AS events is still to be answered. Here, we found downregulation of several splicing factors, including SF3A1, SF3B2, SF3B3, SNRPD4 (SM protein), and U2AF1, which are part of the U2 complex (Figure 1). Their downregulation in resistant cells might suggest alterations in the recognition and usage of the intronic branch site sequence. The downregulation of these factors might drive the AS in resistant cells. In line with this idea, silencing of the splicing factor SF3B1 was shown to reduce the short BRAFV600E isoform in the SKMEL-239 cell line [53]. Note that although SF3B1mutations occur in about 20% of uveal melanomas, the here used SKMEL239 cell line is SF3B1 wild type [54].

As mentioned, we found that the Rho pathway might be regulated by AS in vemurafenib resistant melanoma cells (Figure 6). Different bioinformatic tools identified different AS genes in the Rho pathway, but MPRIP was common to all tools (Figure 6). In the Rho pathway, MPRIP functions as follows. MPRIP binds to MLC phosphatase locating the phosphatase complex to stress fibres thus promoting the dephosphorylation of phosphorylated MLC[31]. It is possible that the here identified AS event in MPRIP impairs this function, meaning that alternatively spliced MPRIP cannot bind and activate MLC phosphatase, thus promoting MLC activity, stress fibre contractility and therapy resistance. It would be interesting to test this hypothesis in future experiments, for example by perturbing MPRIP using RNA-interference or switching the alternative splicing of MPRIP back to normal using splice-switching oligonucleotides [55, 56].

## 5. Conclusions

Alternative splicing of BRAFV600 is a common mechanism for acquired vemurafenib resistance in melanoma. However, the molecular and genetic mechanisms underlying the vemurafenib resistance driven and/or maintained by aberrantly spliced BRAF remains unclear. Deep understanding of the global transcriptional, including alternative splicing, landscape in drug resistant melanoma will be crucial for the development of new therapeutic strategies.

## Supplementary Materials

Figure S1: Western blot of ERK signaling in response to vemurafenib treatment; Figure S2: Relative cell viability measured by MTS assays in response to different doses of vemurafenib treatment for 48 and 72 hours. Figure S3: Principal component analysis (PCA) of RNA-Seq data. Figure S4: Clustergram for DEG; Figure S5: Illustration of the TYR transcripts and domains from the Ensembl genome browser; Figure S6: LeafCutter results for MPRIP; Table S1: Differentially Expressed Genes; Table S2: Differentially expressed spliceosome genes; Table S3: Splicing events; Table S4: BRAF splicing results; Table S5: Overlap of AS analysis for all tools; Table S6: RT-PCR primers sequences; Table S7: IPA comparison of canonical pathway enrichment for AS tools.

## Author Contributions

Conceptualization, A.K., W.K. and H.B.; Methodology, H.B. and A.K.; Formal Analysis, H.B. and A.K.; Investigation, A.K., W.K. and H.B; Writing – Original Draft Preparation, H.B.; Writing – Review & Editing, A.K. and W.K.; Funding Acquisition, W.K.

## Data Availability Statement

Sequencing data are being deposited in ArrayExpress (www.ebi.ac.uk/arrayexpress) and will become available under accession number E-MTAB-11609.

## Acknowledgments

We thank Prof. Poulikos I. Poulikakos (Department of Oncological Sciences Icahn School of Medicine at Mount Sinai, New York, USA) for providing the SK-MEL-239 clone C3 cell line and Dr Jens Rauch for advice with Western blotting.

