## Supplemental figures for "Analysis of alternative mRNA splicing in vemurafenib-resistant melanoma cells"

### Figure S1

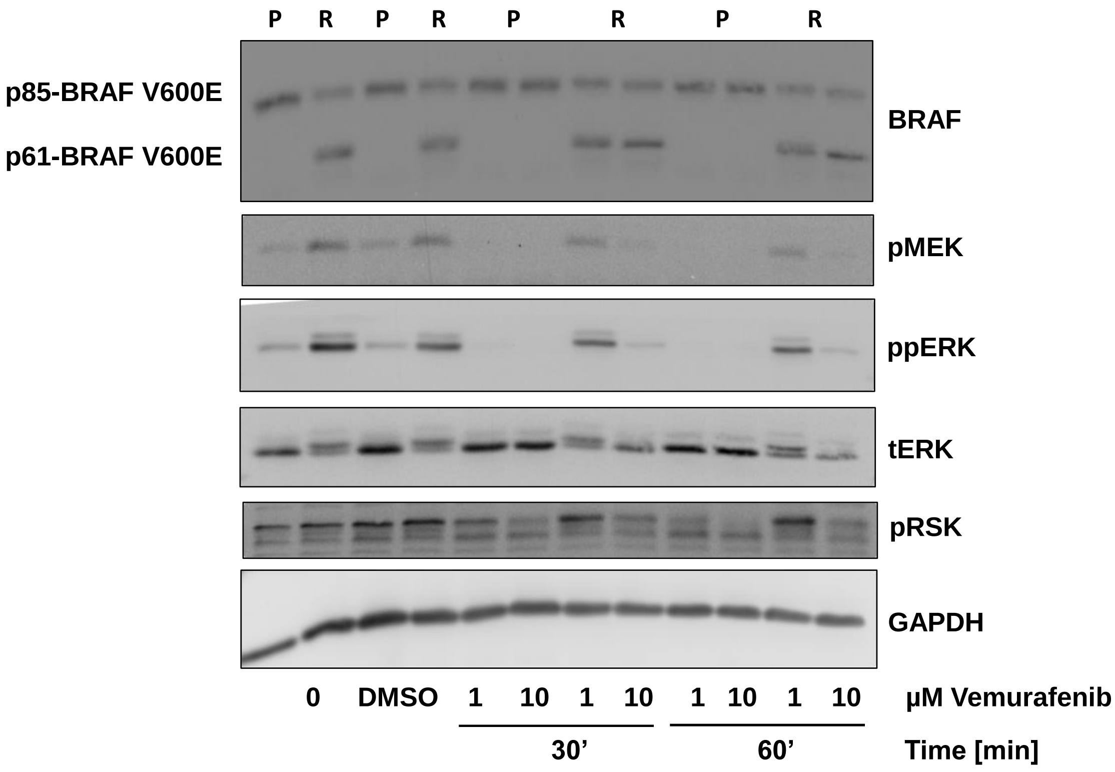


Figure S1. Western blot of ERK signaling in response to vemurafenib treatment. Treatment doses and timepoints in parental (P) and resistant (R) SK-MEL-239 cells are indicated below the image. p85-BRAF indicates full-length BRAF V600E of 85 kDa, and p61 the short splice variant of 61 kDa.

### Figure S2

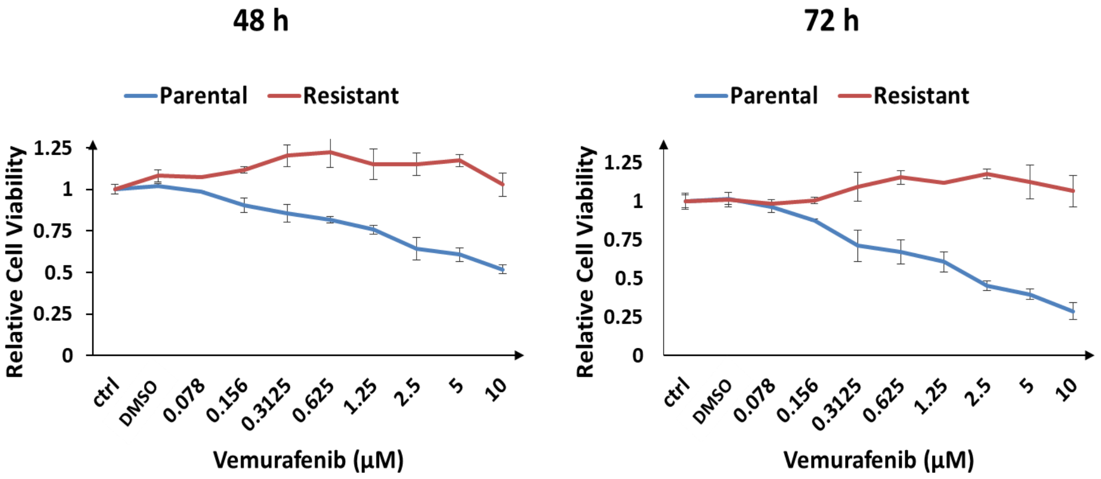


Figure S2. Relative cell viability measured by MTS assays in response to different doses of vemurafenib treatment for 48 and 72 hours, as indicated. ctrl: No treatment, DMSO: DMSO added, numbers: different concentrations of vemurafenib added as indicated.

Figure S3

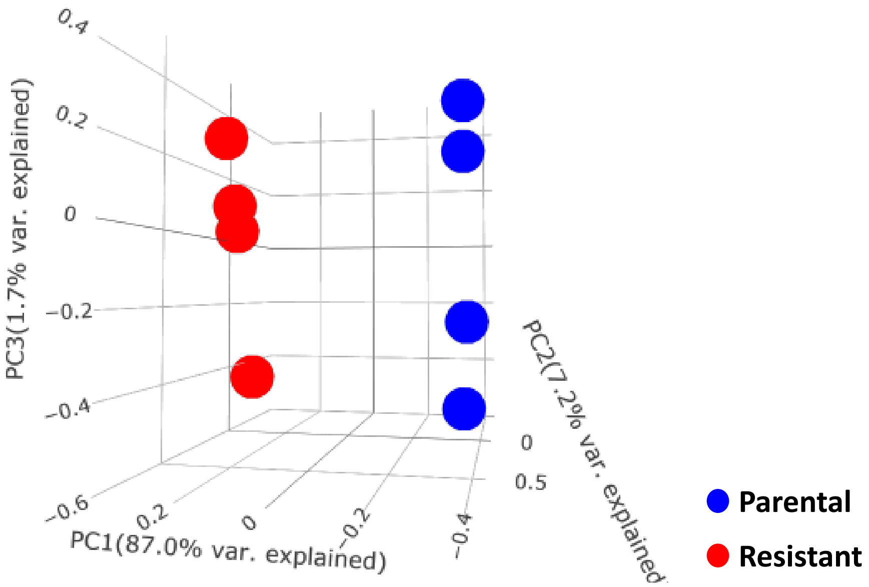


Figure S3. Principal component analysis (PCA) of RNA-Seq data. Four biological replicates in parental (blue circles) and four biological replicates in resistant (red circles) cells using BioJupies. PC1, PC2 and PC3 denote the first, second and third principal component. Indicated percentages show the variance explained by this principal component.

### Figure S4

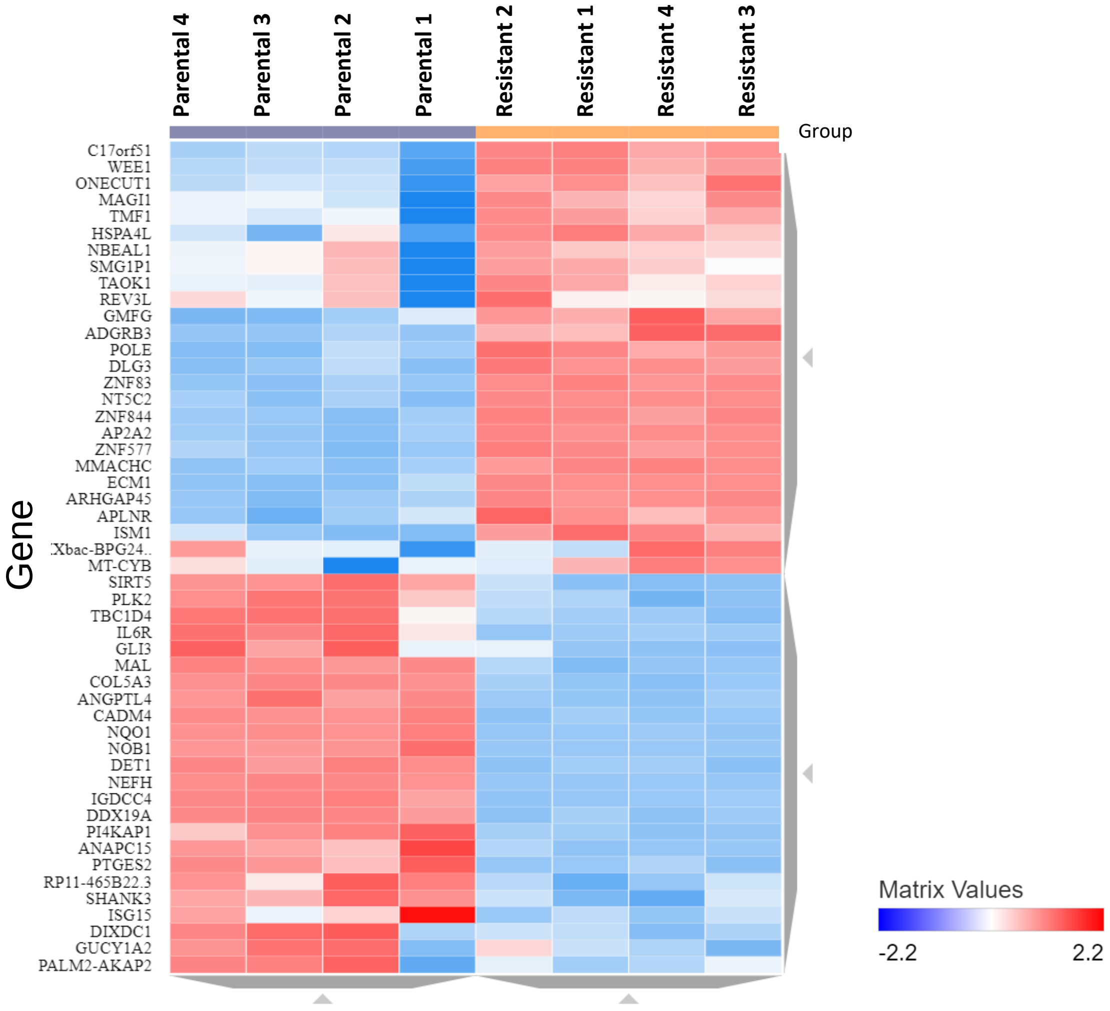


Figure S4. Clustergram for DEG. Top 50 variable genes (rows) for four biological replicates in parental and resistant cells (column) were obtained using Clustergrammer (Fernandez et al., 2017) in BioJupies. Colors represent normalized gene expression values.

### Figure S5

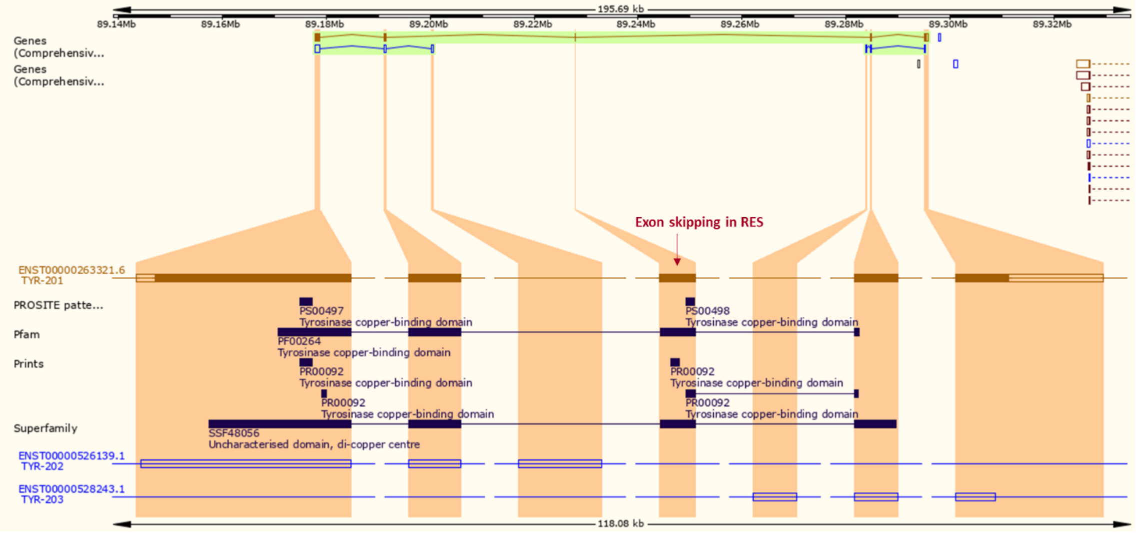


Figure S5. Illustration of the TYR transcripts and domains from the Ensembl genome browser. The blue arrow shows which exon is skipped. The skipped exon contains a copper binding domain.

### Figure S6

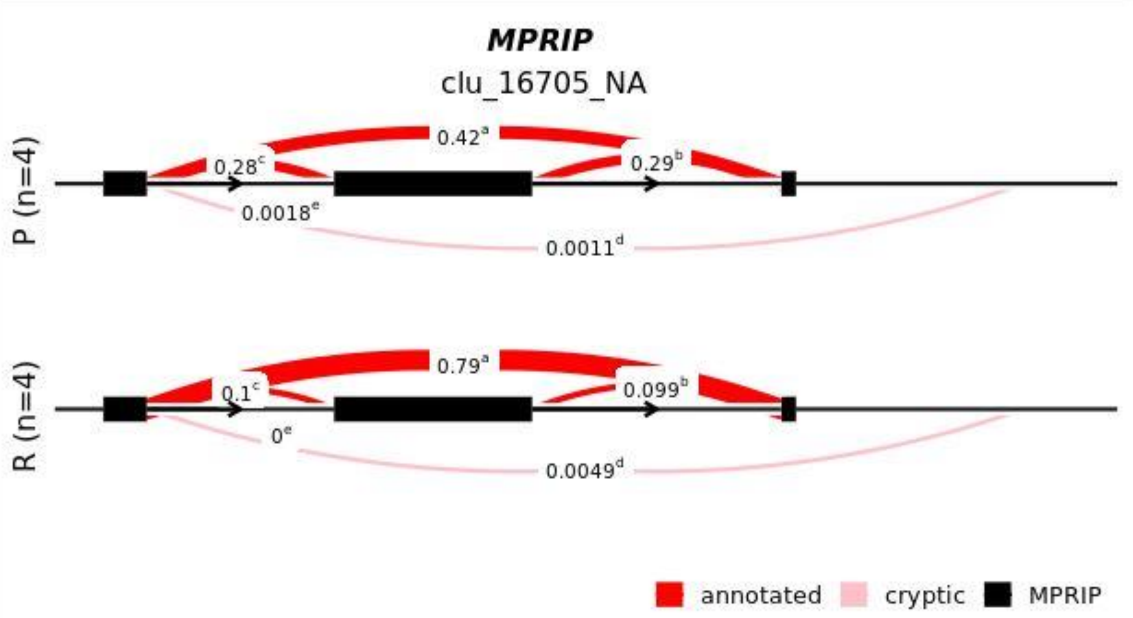


| | chr | start | end | verdict | ΔPSI |
|---|---|---|---|---|---|
| a | chr17 | 17180088 | 17184823 | annotated | 0.368 |
| b | chr17 | 17180669 | 17184823 | annotated | -0.184 |
| c | chr17 | 17180088 | 17180607 | annotated | -0.182 |
| d | chr17 | 17180088 | 17188328 | cryptic_threeprime | -0.002 |

Figure S6. LeafCutter results for MPRIP. Top panel, cluster plot for the comparison of the junction usage in parental versus resistant cells. Numbers are PSI (percentage spliced in) values. The table below shows ΔPSI for each junction usage.
