## Supplementary figures and images for "Analysis of alternative mRNA splicing in vemurafenib-resistant melanoma cells"

### Table S6

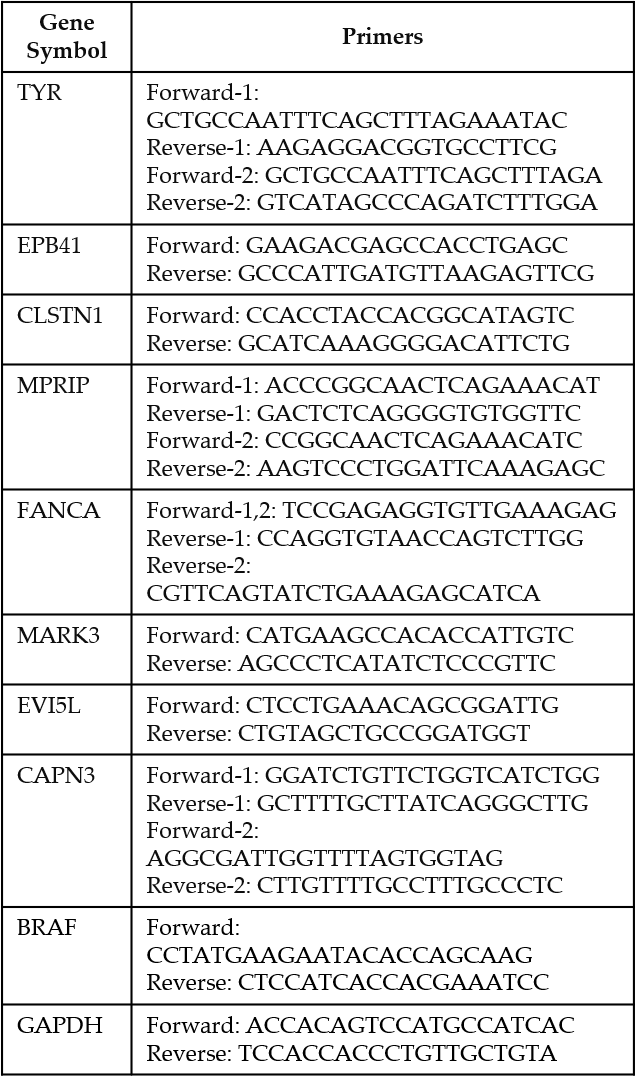

Table S6

Table S6. RT-PCR primers sequences.
